# Cluster Analysis Based on Fasting and Postprandial Plasma Glucose and Insulin Concentrations

**DOI:** 10.1101/861468

**Authors:** Miguel Altuve, Erika Severeyn

## Abstract

Plasma glucose and insulin concentrations are clinical markers used in the diagnosis of metabolic diseases, particularly prediabetes and diabetes. In this paper, we carried out a cluster analysis using plasma glucose and insulin data in fasting and two-hour postprandial. Different clustering experiments were performed by changing the attributes, from one (fasting glucose) to four (fasting and postprandial glucose and insulin) attribute input to a k-means clustering algorithm. Based on the elbow and silhouette methods, three clusters were chosen to carry out the clustering experiments. The Pearson correlation coefficient was used to assess the dependence between the glucose and insulin levels for each cluster created. Results show that one cluster contained prediabetics, another cluster contained diabetics, and subjects without prediabetes and diabetes were assigned to another cluster. Although age was not used as an attribute, we have found that subjects in the three clusters have a different age range. Finally, significant correlations were found between insulin levels in fasting and postprandial and between glucose levels in fasting and postprandial. These associations were stronger in the cluster containing diabetics, where insulin production or action is compromised.

## 1 Introduction

Diabetes is a progressive disease characterized by high blood glucose levels due to poor performance in the production or action of insulin [6, 17]. There is no clear knowledge about the instant in which diabetes occurs but there are pre-diabetic conditions, such as impaired glucose tolerance (IGT) and impaired fasting glucose (IFG), that predispose the development of diabetes [10] and increase the risk of cardiovascular diseases [19].

The increase of the population in urban areas and the proliferation of a sedentary lifestyle have influenced the prevalence diabetes and prediabetes worldwide. For instance, two thirds of adults with diabetes live in urban settings (298 million people) whereas a third lives in rural areas (153 million people) [12]. The prevalence of diabetes in adults in 2017 was estimated in 425 million cases (almost 9% of adults) but it is estimated that 629 million people (about 10% of adults) will suffer from diabetes in 2045. Similarly, the prevalence of IGT in adults in 2017 was estimated in 374 million cases (almost 8% of adults), 69% of which live in low and middle income countries, but it is estimated that 587 million adults (8.4% of adults) will suffer from IGT in 2045 [12]. The North American and Caribbean region has the highest prevalence of IGT (13.6%) and the Southeast Asia region has the lowest one (3.4%). The prevalence of IFG is between 43.9% and 58% for Caucasians and between 29.2% and 48.1% for Asians [28].

Diabetes, IGT and IFG can be diagnosed from the oral glucose tolerance test (POTG) [20], a clinical test consisting of five plasma glucose and insulin measurements, one in fasting and, after oral intake of 75 grams of liquid anhydrous glucose, four others each 30 minutes. IGT is diagnosed when plasma glucose levels at 120 minutes are between 140 and 200 mg/dL and IFG is diagnosed when fasting glucose levels are between 100 and 125 mg/dL [6].

Numerous methodologies have been used for the diagnosis of diabetes and prediabetes. Support vector machines, genetic algorithms and k-means have been used for the diagnosis of diabetes [25] and gestational diabetes [23] using databases that include plasma glucose, plasma insulin, blood pressure, among others. Likewise, linear regressions has been used for the diagnosis of IGT to correlate plasma metabolite values with insulin and OGTT glucose [26]. Additionally, machine learning algorithms have been explored to predict the evolution towards diabetes in patients at high risk [11].

Diabetes is a multifactorial disease that usually comes in concomitance with other pathologies, such as insulin resistance, hypertension, dyslipidemia, metabolic syndrome, among others. Some studies have explored the characterization of diabetics in more specific groups with particular characteristics, the results of these investigations have revealed that a more specific characterization of diabetic patients could improve the quality of life of diabetic patients by designing treatments more appropriate to their type of diabetes and even avoid long-term diabetic complications such as cardiovascular problems [1, 4].

Fasting and postprandial blood glucose and insulin levels defined in the literature are used to identify prediabetic and diabetic conditions. In a recent study, we have used these ranges of values and combined them to better characterize the metabolic conditions of individuals [4]. Specifically, we define 28 different classes, of which four correspond to the normal condition, twelve to the condition of prediabetes and twelve to the condition of diabetes. In this work, we propose to cluster these subjects using fasting and postprandial glucose and insulin values in order to find characteristic patterns of glucose and insulin that reveal important information about the processes of deterioration of the metabolic condition. In this sense, we propose to divide the data set into three groups using the k-means algorithm, which allows to find natural groups in the data set by means of a similarity measure based on the distance between individuals, in such a way that the individuals within the same cluster are similar to each other and share the same attributes (characteristics) and these in turn are different from the individuals of the other groups [9]. The advantage of clustering is that it allows the subjects to be grouped without having a priori knowledge of how to do it (it is a type of unsupervised learning) since the data is not labeled [24].

Prediabetic conditions can be reversed with simple changes in people’s lifestyle [29] but diabetes can only be controlled with medical treatment. Therefore, early diagnosis of prediabetic conditions is of paramount importance in the prevention of diabetes [5]. The OGTT is an invasive and expensive test, the application of methodologies that improve the diagnosis of diabetes and prediabetes by decreasing the amount of glucose and insulin measurements and including non-invasive variables such as age could be of great help especially in developing countries [13]. There is evidence that early diagnosis and prediction of the onset of diabetes are vital for the delay of disease progression, adequate selection of the treatment (personalized medicine), prolongation of life expectancy, symptom relief and prevention of related complications [21].

## 2 Methods

### 2.1 Database

The age and levels of glucose and insulin in plasma in fasting from 2835 women, collected in a previous study at Caracas University Hospital, Venezuela, were employed in this work. Please refer to [4] for a more detailed explanation of the clinical protocol and the database in general. The database is freely available at [2].

### 2.2 k-means clustering

In this work, we focused on analyzing the result of the clustering for different combinations of attributes. Specifically, we performed four different clustering experiments by changing the attributes as follows:

1. The first clustering experiment was performed using only fasting glucose levels (*G*_0_). This experiment is important since estimating the fasting glucose concentration is a feasible task to be performed by an inexperienced person using, for example, commercial devices, such as a glucometer or a smartphone application [8, 27], or wearable technology, such as a contact lens [16, 7]. Therefore, discovering physiological patterns or clinical indicators using only fasting blood glucose concentration could have a significant impact on society since people could obtain useful information without going to a specialized clinical laboratory.
2. The second clustering experiment uses fasting and postprandial glucose levels (*G*_0_&*G*_120_). Although two blood samples are required to perform this experiment and ingest a certain glucose concentration, which indicates that it is a more expensive and complicated procedure than the previous one, most specialized laboratories perform this type of test to diagnose metabolic diseases. Therefore, discovering physiological patterns or clinical indicators using fasting and postprandial blood glucose concentrations could be a useful tool for clinical laboratories.
3. The third one uses fasting glucose and insulin levels (*G*_0_&*I*_0_). The measurement of the concentration of insulin in the blood is carried out with specialized equipment in a clinical laboratory, therefore, it is an expensive procedure that requires specialized personnel. This experiment, like the previous one, could yield useful information for clinical laboratories.
4. The fourth and final clustering experiment uses fasting and postprandial glucose and insulin levels (*G*_0_&*G*_120_&*I*_0_&*I*_120_). Observing these four variables at the same time can help contribute to discovering patterns or clinical indicators in the data. Besides, we recall that the database was classified into 28 classes based on fasting and postprandial glucose and insulin values defined in the literature [4]. However, this test would require two blood samples, the intake of a certain amount of glucose, and a specialized instrument to determine the concentration of insulin in blood (glucose concentration can also be obtained with a specialized instrument, however, as mentioned, a person can estimate its blood glucose concentration using, for example, a glucometer). This experiment would, therefore, have an impact on clinical laboratories.

Since our objective is to group the subjects based on blood glucose and insulin concentrations, the age variable was not considered as an attribute in the clustering process, however, it was used for analyzing the groups created. The number of subjects per group was determined for each clustering experiment.

The squared Euclidean distance was used as the distance measure in which, for each attribute, each centroid corresponds to the mean of the values of the attribute of the individuals assigned to that cluster.

Since hyperparameter *k* must be specified prior to the clustering process, the elbow and silhouette methods were used to estimate the number of clusters. In the case of the elbow method, for different values of *k*, we performed the clustering process and compute the percentage of variance explained. Then, the optimal number of clusters corresponds to the lowest *k* that gives 90% of the percentage of variance explained. The elbow method by considering the total within cluster sum of squares (WCSS) as a function of the number of clusters was also considered. In the case of the Silhouette clustering evaluation criterion, again for different values of *k*, we performed the clustering process and computed the average Silhouette of observations. Then, the optimal number of clusters is the one that provides the highest average Silhouette value.

### 2.3 Statistical analysis

We computed the mean and standard deviation of the variables age, and fasting and postprandial glucose and insulin levels of the subjects assigned to the clusters. The Kruskal-Wallis nonparametric statistical test was performed to find significant differences in the variables age, and fasting and postprandial glucose and insulin levels between clusters (independent samples) followed by the Tukey’s honestly significant difference test as a post hoc test. The Wilcoxon signed rank test was used to assess the significant differences in the variables glucose and insulin between fasting and posprandial (dependent samples). The Pearson correlation coefficient was used to assess the linear dependence between the glucose and insulin levels per cluster. A *p*-value less than or equal to 5% was considered to be statistically significant.

## 3 Results

A capsule in Code Ocean to reproduce the results of this paper is available at [3]. Figure 1 shows the results of applying the elbow and average Silhouette methods for deciding the number of clusters. Based on the results, we decided to use three clusters for all experiments.

**Figure 1:**
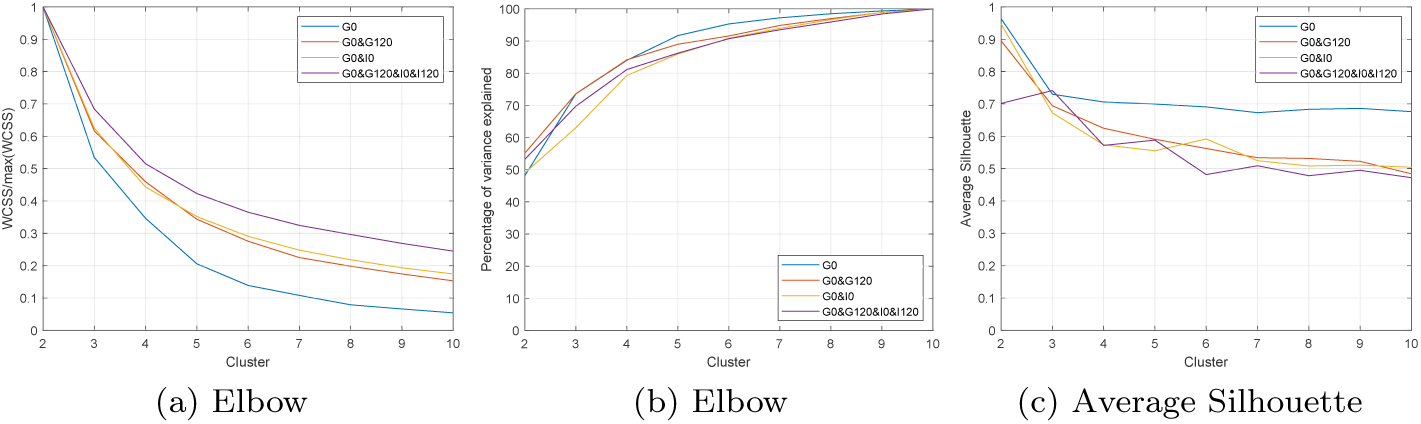
Elbow and average Silhouette methods for different combination of the attributes.

Table 1 shows the number of subjects assigned to each cluster for the different clustering experiments carried out, i.e. by considering *G*_0_, *G*_0_&*G*_120_, *G*_0_&*I*_0_ and *G*_0_&*G*_120_&*I*_0_&*I*_120_ as attributes of the clustering process.

**Table 1:** Number of subjects per cluster for different attributes used by the clustering algorithm.

| Cluster | Attributes |  |  |  |
| --- | --- | --- | --- | --- |
| | $G_0$ | $G_0 \& G_{120}$ | $G_0 \& I_0$ | $G_0 \& I_0 \& G_{120} \& I_{120}$ |
| C1 | 2080 | 2013 | 2124 | 2256 |
| C2 | 724 | 724 | 680 | 441 |
| C3 | 31 | 98 | 31 | 138 |

Table 2 shows the result of the variables age, *G*_0_, *G*_120_, *I*_0_ and *G*_120_ of the subjects assigned to each cluster for each clustering experiment carried out. Significant differences of the variables between clusters and of the cluster between the levels of glucose and insulin in fasting and postprandial are also shown.

**Table 2:** Mean ± standard deviation of the variables per cluster for each attribute used in the clustering process.

| Variable | Cluster | Attributes |  |  |  |
| --- | --- | --- | --- | --- | --- |
|  |  | G0 | G0&G120 | G0&I0 | G0&I0&G120&I120 |
| Age | C1 | 39.83 ± 14.57 <sup>a,b</sup> | 40.05 ± 14.74 <sup>a,b,c</sup> | 40.07 ± 14.61 <sup>a,b</sup> | 41.87 ± 14.81 <sup>b,c</sup> |
| | C2 | 50.11 ± 12.96 | 48.33 ± 13.46 | 50.04 ± 13.00 | 43.07 ± 15.06 |
| | C3 | 50.65 ± 10.04 | 52.00 ± 10.67 | 50.65 ± 10.04 | 52.58 ± 10.43 |
| G0 | C1 | 91.16 ± 7.25 <sup>a,b,c,$\alpha$</sup> | 93.13 ± 9.50 <sup>a,b,c,$\alpha$</sup> | 91.44 ± 7.44 <sup>a,b,c,$\alpha$</sup> | 95.73 ± 12.38 <sup>a,b,c,$\alpha$</sup> |
| | C2 | 116.09 ± 13.24 <sup>$\alpha$</sup> | 107.01 ± 16.52 <sup>$\alpha$</sup> | 116.82 ± 13.35 <sup>$\alpha$</sup> | 99.37 ± 12.83 <sup>$\alpha$</sup> |
| | C3 | 234.48 ± 64.54 <sup>$\alpha$</sup> | 163.22 ± 61.80 <sup>$\alpha$</sup> | 234.48 ± 64.54 <sup>$\alpha$</sup> | 153.25 ± 55.44 <sup>$\alpha$</sup> |
| G120 | C1 | 107.54 ± 23.62 <sup>a,b,c</sup> | 100.52 ± 13.53 <sup>a,b,c</sup> | 107.88 ± 23.95 <sup>a,b,c</sup> | 107.52 ± 21.75 <sup>a,b,c</sup> |
| | C2 | 145.30 ± 50.34 | 149.25 ± 22.89 | 146.70 ± 50.96 | 136.33 ± 29.80 |
| | C3 | 313.55 ± 112.95 | 287.78 ± 66.85 | 313.55 ± 112.95 | 260.34 ± 70.39 |
| I0 | C1 | 8.01 ± 6.70 <sup>a,b,$\beta$</sup> | 8.03 ± 6.74 <sup>a,b,c,$\beta$</sup> | 7.81 ± 6.48 <sup>a,b,$\beta$</sup> | 7.35 ± 5.82 <sup>a,b,c,$\beta$</sup> |
| | C2 | 11.01 ± 8.79 <sup>$\beta$</sup> | 10.21 ± 8.00 <sup>$\beta$</sup> | 11.82 ± 9.13 <sup>$\beta$</sup> | 15.13 ± 9.80 <sup>$\beta$</sup> |
| | C3 | 12.39 ± 8.87 <sup>$\beta$</sup> | 14.99 ± 11.45 <sup>$\beta$</sup> | 12.39 ± 8.87 <sup>$\beta$</sup> | 12.82 ± 9.96 <sup>$\beta$</sup> |
| I120 | C1 | 62.94 ± 46.03 <sup>a,b,c</sup> | 55.08 ± 38.74 <sup>a,b,c</sup> | 62.08 ± 44.95 <sup>a,b,c</sup> | 47.79 ± 22.06 <sup>a,b,c</sup> |
| | C2 | 73.27 ± 51.83 | 91.35 ± 56.91 | 76.61 ± 54.37 | 153.11 ± 48.34 |
| | C3 | 38.45 ± 28.57 | 83.03 ± 60.84 | 38.45 ± 28.57 | 71.15 ± 41.02 |

<sup>a</sup> The variable shows significant difference between clusters C1 and C2.
<sup>b</sup> The variable shows significant difference between clusters C1 and C3.
<sup>c</sup> The variable shows significant difference between clusters C2 and C3.
<sup>$\alpha$</sup> The cluster shows significant difference between variables G0 and G120.
<sup>$\beta$</sup> The cluster shows significant difference between variables I0 and I120.

Figure 2 shows box plot of the variables per cluster for each attribute combination of the clustering process.

**Figure 2:**
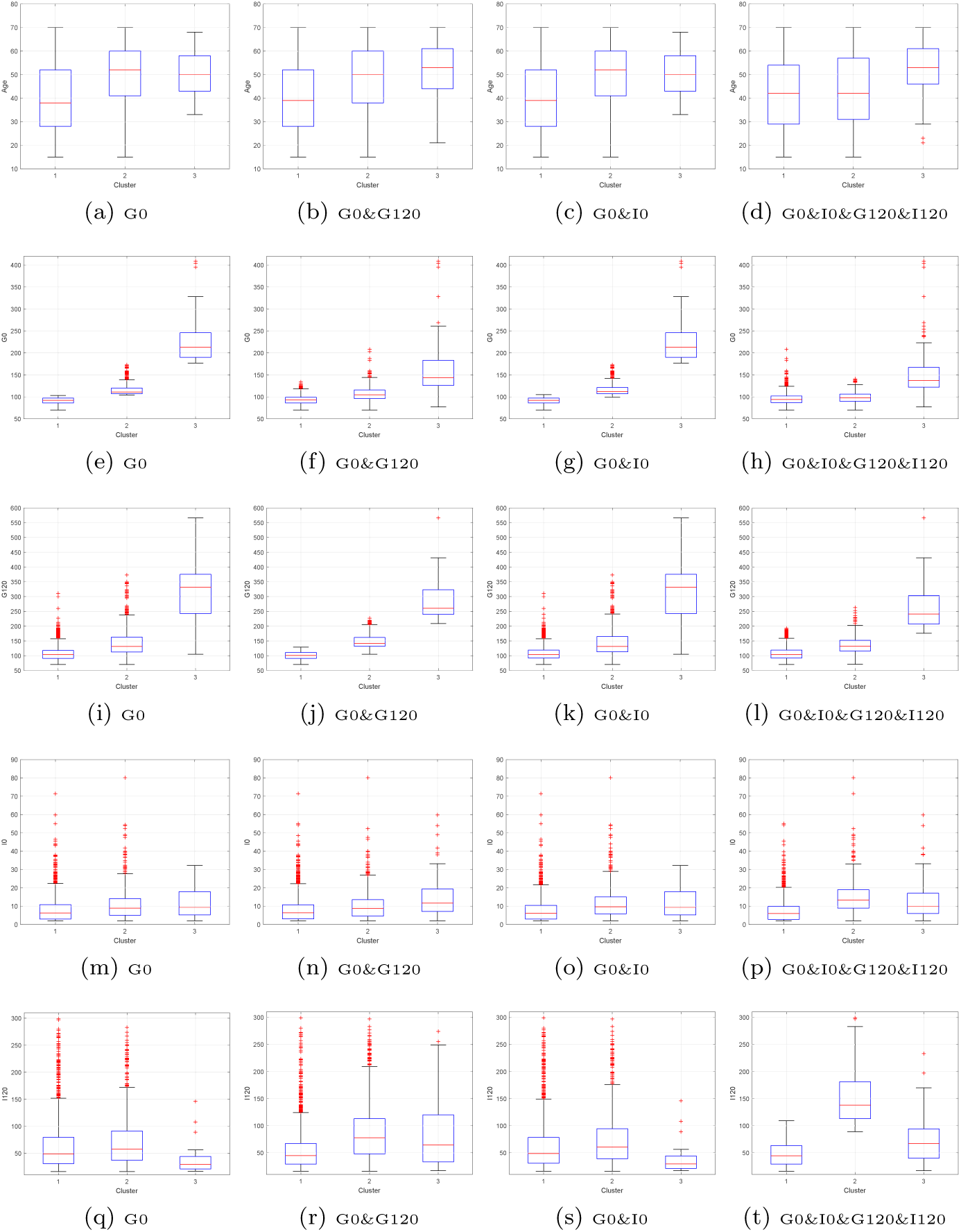
Box plots of the variables (rows) per cluster for each attribute combination (columns) used in the clustering process.

Table 3 shows the correlations between the variables of glucose and insulin in fasting and postprandial per cluster.

**Table 3:** Correlations between variables per cluster. Correlations statistically significant and greater than |±0.5| are shown in bold text.

| Cluster | Variable | Attributes |  |  |  |  |  |  |  |  |  |  |  |
| --- | --- | --- | --- | --- | --- | --- | --- | --- | --- | --- | --- | --- | --- |
|  |  | G0 |  |  | G0&G120 |  |  | G0&I0 |  |  | G0&I0&G120&I120 |  |  |
|  |  | G120 | I0 | I120 | G120 | I0 | I120 | G120 | I0 | I120 | G120 | I0 | I120 |
| C1 | G0 | 0.251 | 0.138 | 0.092 | 0.259 | 0.159 | -0.008 | 0.262 | 0.088 | 0.060 | 0.438 | 0.163 | 0.027 |
|  | G120 |  | 0.142 | 0.448 |  | 0.089 | 0.338 |  | 0.121 | 0.432 |  | 0.016 | 0.289 |
|  | I0 |  |  | **0.513** |  |  | 0.492 |  |  | **0.504** |  |  | 0.411 |
| C2 | G0 | **0.587** | 0.112 | 0.038 | 0.196 | 0.117 | -0.165 | **0.588** | 0.024 | -0.023 | 0.430 | 0.155 | 0.049 |
|  | G120 |  | 0.095 | 0.267 |  | -0.019 | 0.110 |  | 0.056 | 0.239 |  | -0.049 | 0.151 |
|  | I0 |  |  | 0.492 |  |  | 0.484 |  |  | 0.492 |  |  | 0.282 |
| C3 | G0 | **0.714** | 0.319 | -0.030 | **0.701** | -0.058 | -0.439 | **0.714** | 0.319 | -0.030 | **0.699** | 0.093 | -0.361 |
|  | G120 |  | 0.345 | -0.089 |  | -0.084 | -0.377 |  | 0.345 | -0.089 |  | 0.074 | -0.231 |
|  | I0 |  |  | **0.530** |  |  | **0.559** |  |  | **0.530** |  |  | 0.460 |

## 4 Discussion

In all the experiments carried out with the different attribute combinations, subjects in cluster C1 have G0 and G120 lower than subjects in cluster C2, and they, in turn, have G0 and G120 lower than subjects in cluster C3. It is important to note that we have named cluster C1 the one with the most subjects and cluster C3 the one with fewer subjects, as shown in Table 1. Moreover, the results of the clustering algorithm are in accordance with the diagnostic values of diabetes and prediabetes stated by the American Diabetes Association [6]: cluster C2 contains prediabetics, cluster C3 contains diabetic, and cluster C1 contains subjects without any of the above pathologies.

Age is an important factor in the development of diabetes, and this is reflected in the clustering performed. In all experiments, subjects in cluster C1 were younger than subjects in cluster C2, and they, in turn, were younger than subjects in cluster C3. Consequently, as we age, we are more likely to suffer from diabetes. Moreover, a chronological trend was observed: the youngest tends to be healthy (cluster C1), prediabetes tends to occur at middle-aged adults (cluster C2) and diabetes tends to appear at an old-age (cluster C3) [18].

By considering the insulin concentration, on the one hand, in all the experiments, cluster C1 grouped subjects with lower I0 and greater insulin sensitivity (mean HOMAIR < 1.84). This is an expected result since these subjects tend to have normal fasting glucose and insulin levels. Clustering experiments with G0, G0&G120, and G0&I0 as attributes, grouped subjects with higher I0 levels in cluster C3. These subjects have lower insulin sensitivity (mean HOMA-IR*>* 6.04). It is known that low insulin sensitivity can cause the development of diabetes. This can be reflected by the fact that subjects in cluster C2 have a mean HOMA-IR*>* 2.6, indicating that these subjects (with glucose values consistent with prediabetes) have insulin resistance [14]. On the other hand, in all the experiments, cluster C2 contains subjects with elevated levels of I120, which could indicate that subjects with prediabetes tend to produce more insulin for glucose metabolism than normal. Moreover, cluster C3 contains the lowest level of I120 when the clustering is carried out with G0 and G0&I0 as attributes. Subjects in cluster C3 are mostly diabetic as indicated by their fasting and postprandial glucose levels, their insulin production is thus compromised due to the diabetic condition [15].

It is interestingly to note the behavior of the correlations between the variables as the metabolic disease progresses (from cluster C1 to C3). For instance, clustering experiments with G0 and G0&I0 show significant correlations above 0.5 between fasting and postprandial insulin in cluster C1 and between fasting and postprandial glucose in cluster C2. The association between fasting and postprandial glucose appears in cluster C2 (mostly composed by prediabetics) but it is then strengthened in cluster C3 (mostly composed by diabetics), with correlations around 0.7, for all the clustering experiments. This suggests that the association between fasting and postprandial plasma glucose concentration is marked in subjects with problems with insulin production or action [22].

## 5 Conclusions

This work has shown that the k-means clustering algorithm with three clusters groups the subjects into healthy, prediabetic and diabetic, by using fasting and postprandial glucose and insulin levels. In addition, it also groups subjects with low insulin sensitivity. Age can also be considered an important factor in the development of diabetes since subjects with diabetes and prediabetes (clusters C3 and C2, respectively) were among the older subjects. Low insulin sensitivity could be another risk factor in the development of diabetes. Clusters C2 and C3 contain subjects with HOMA-IR*>* 2.5, being the subjects of cluster C3 (diabetics) those who presented a greater detriment in insulin sensitivity.

## Author Contributions

Miguel Altuve carried out the experiment and contributed to the conception of the study, and the writing and revising of the manuscript. Erika Severeyn contributed to the conception of the study, the analysis of the results and the writing and revising of the manuscript.

## References

[1] E. Ahlqvist, P. Storm, A. Käräjämäki, M. Martinell, M. Dorkhan, A. Carlsson, P. Vikman, R. B. Prasad, D. M. Aly, P. Almgren, et al. Novel subgroups of adult-onset diabetes and their association with outcomes: a data-driven cluster analysis of six variables. The lancet Diabetes & endocrinology, 6(5):361–369, 2018. ISSN 2213-8587. doi: 10.1016/S2213-8587(18)30051-2.

[2] M. Altuve and E. Severeyn. Fasting and postprandial glucose and insulin dataset. http://dx.doi.org/10.21227/5g52-jc59, 2019.

[3] M. Altuve and E. Severeyn. Cluster analysis based on fasting and postprandial plasma glucose and insulin concentrations [source code]. https://doi.org/10.24433/CO.7408455.v1, 11 2019.

[4] M. Altuve and E. Severeyn. Joint analysis of fasting and postprandial plasma glucose and insulin concentrations in venezuelan women. Diabetes & Metabolic Syndrome: Clinical Research & Reviews, 13(3):2242–2248, 2019. ISSN 1871-4021. doi: 10.1016/j.dsx.2019.05.029.

[5] R. Ambady and S. Chamukuttan. Early diagnosis and prevention of diabetes in developing countries. Reviews in Endocrine and Metabolic Disorders, 9(3): 193–201, 2008. ISSN 1573-2606. doi: 10.1007/s11154-008-9079-z.

[6] American Diabetes Association. 2. classification and diagnosis of diabetes: Standards of medical care in diabetes—2019. Diabetes Care, 42(Supplement 1):S13–S28, 2019. ISSN 0149-5992. doi: 10.2337/dc19-S002.

[7] R. Badugu, E. A. Reece, and J. R. Lakowicz. Glucose-sensitive silicone hydrogel contact lens toward tear glucose monitoring. Journal of biomedical optics, 23(5): 057005, 2018. ISSN 1083-3668. doi: 10.1117/1.JBO.23.5.057005.

[8] A. J. Bandodkar, S. Imani, R. Nunez-Flores, R. Kumar, C. Wang, A. V. Mohan, J. Wang, and P. P. Mercier. Re-usable electrochemical glucose sensors integrated into a smartphone platform. Biosensors and Bioelectronics, 101:181–187, 2018. ISSN 0956-5663. doi: 10.1016/j.bios.2017.10.019.

[9] S. Bandyopadhyay and S. Saha. Unsupervised classification: similarity measures, classical and metaheuristic approaches, and applications. Springer Science & Business Media, 2012.

[10] N. Bansal. Prediabetes diagnosis and treatment: A review. World journal of diabetes, 6(2):296–303, 2015. ISSN 1948-9358. doi: 10.4239/wjd.v6.i2.296.

[11] A. Cahn, A. Shoshan, T. Sagiv, R. Yesharim, I. Raz, and R. Goshen. Use of a machine learning algorithm improves prediction of progression to diabetes, 2018. ISSN 0012-1797.

[12] N. Cho, J. Shaw, S. Karuranga, Y. Huang, J. da Rocha Fernandes, A. Ohlrogge, and B. Malanda. IDF diabetes atlas: Global estimates of diabetes prevalence for 2017 and projections for 2045. Diabetes research and clinical practice, 138: 271–281, 2018. ISSN 0168-8227. doi: 10.1016/j.diabres.2018.02.023.

[13] C. K. Chow, C. Ramasundarahettige, W. Hu, K. F. AlHabib, A. Avezum Jr, X. Cheng, J. Chifamba, G. Dagenais, A. Dans, B. A. Egbujie, et al. Availability and affordability of essential medicines for diabetes across high-income, middle-income, and low-income countries: a prospective epidemiological study. The lancet Diabetes & endocrinology, 6(10):798–808, 2018. ISSN 2213-8587. doi: 10.1016/S2213-8587(18)30233-X.

[14] M. P. Czech. Insulin action and resistance in obesity and type 2 diabetes. Nature medicine, 23(7):804–814, 2017. ISSN 1078-8956. doi: 10.1038/nm.4350.

[15] R. A. DeFronzo, E. Ferrannini, L. Groop, R. R. Henry, W. H. Herman, J. J. Holst, F. B. Hu, C. R. Kahn, I. Raz, G. I. Shulman, et al. Type 2 diabetes mellitus. Nature reviews Disease primers, 1:15019, 2015. ISSN 2056-676X. doi: 10.1038/nrdp.2015.19.

[16] N. M. Farandos, A. K. Yetisen, M. J. Monteiro, C. R. Lowe, and S. H. Yun. Contact lens sensors in ocular diagnostics. Advanced healthcare materials, 4(6): 792–810, 2015. ISSN 2192-2640. doi: 10.1002/adhm.201400504.

[17] V. A. Fonseca. Defining and characterizing the progression of type 2 diabetes. Diabetes care, 32(suppl 2):S151–S156, 2009. ISSN 0149-5992. doi: 10.2337/dc09-S301.

[18] K. H. Ha and D. J. Kim. Trends in the diabetes epidemic in korea. Endocrinology and Metabolism, 30(2):142–146, 2015. ISSN 2093-596X. doi: 10.3803/EnM.2015.30.2.142.

[19] Y. Huang, X. Cai, W. Mai, M. Li, and Y. Hu. Association between prediabetes and risk of cardiovascular disease and all cause mortality: systematic review and meta-analysis. Bmj, 355:i5953, 2016. ISSN 0959-8138. doi: 10.1136/bmj.i5953.

[20] A. Hulman, D. Vistisen, C. Glümer, M. Bergman, D. R. Witte, and K. Færch. Glucose patterns during an oral glucose tolerance test and associations with future diabetes, cardiovascular disease and all-cause mortality rate. Diabetologia, 61(1): 101–107, 2018. ISSN 1432-0428. doi: 10.1007/s00125-017-4468-z.

[21] I. Kavakiotis, O. Tsave, A. Salifoglou, N. Maglaveras, I. Vlahavas, and I. Chouvarda. Machine learning and data mining methods in diabetes research. Computational and structural biotechnology journal, 15:104–116, 2017. ISSN 2001-0370. doi: 10.1016/j.csbj.2016.12.005.

[22] E. B. Ketema and K. T. Kibret. Correlation of fasting and postprandial plasma glucose with hba1c in assessing glycemic control; systematic review and meta-analysis. Archives of Public Health, 73(1):43, 2015. ISSN 2049-3258. doi: 10.1186/s13690-015-0088-6.

[23] T. Santhanam and M. Padmavathi. Application of K-means and genetic algorithms for dimension reduction by integrating SVM for diabetes diagnosis. Procedia Computer Science, 47:76–83, 2015. ISSN 1877-0509. doi: 10.1016/j.procs.2015.03.185.

[24] A. Saxena, M. Prasad, A. Gupta, N. Bharill, O. P. Patel, A. Tiwari, M. J. Er, W. Ding, and C.-T. Lin. A review of clustering techniques and developments. Neurocomputing, 267:664–681, 2017. ISSN 0925-2312. doi: 10.1016/j.neucom.2017.06.053.

[25] P. M. Shakeel, S. Baskar, V. S. Dhulipala, and M. M. Jaber. Cloud based framework for diagnosis of diabetes mellitus using K-means clustering. Health information science and systems, 6(1):16, 2018. ISSN 2047-2501. doi: 10.1007/s13755-018-0054-0.

[26] C. Wildberg, A. Masuch, K. Budde, G. Kastenmüller, A. Artati, W. Rathmann, J. Adamski, T. Kocher, H. Völzke, M. Nauck, et al. Plasma metabolomics to identify and stratify patients with impaired glucose tolerance. The Journal of Clinical Endocrinology & Metabolism, 2019. ISSN 0021-972X. doi: 10.1210/jc.2019-01104.

[27] J. Yang. Blood glucose monitoring with smartphone as glucometer. Electrophoresis, 40(8):1144–1147, 2019. ISSN 0173-0835. doi: 10.1002/elps.201800295.

[28] W. Yip, I. Sequeira, L. Plank, and S. Poppitt. Prevalence of pre-diabetes across ethnicities: a review of impaired fasting glucose (IFG) and impaired glucose tolerance (IGT) for classification of dysglycaemia. Nutrients, 9(11):1273, 2017. ISSN 2072-6643. doi: 10.3390/nu9111273.

[29] P. Zimmet, K. Alberti, and J. Shaw. Global and societal implications of the diabetes epidemic. Nature, 414(6865):782–787, 2001. ISSN 0028-0836. doi: 10.1038/414782a.

